# ChIPflow: from raw data to epigenomic dynamics

**DOI:** 10.1101/2021.02.02.429342

**Authors:** Maëlle Daunesse, Rachel Legendre, Hugo Varet, Adrien Pain, Claudia Chica

## Abstract

We present ChIPflow, a Snakemake-based pipeline for epigenomic data from the raw fastq files to the differential analysis. It can be applied to any chromatin factor, e.g. histone modification or transcription factor, which can be profiled with ChIP-seq. ChIPflow streamlines critical steps like the quality assessment of the immunoprecipitation using cross-correlation and the replicate comparison for both narrow and broad peaks. For the differential analysis ChIPflow provides linear and nonlinear methods for normalisation between samples as well as conservative and stringent models for estimating the variance and testing the significance of the observed binding/marking differences.

ChIPflow can process in parallel multiple chromatin factors with different experimental designs, number of biological replicates and/or conditions. It also facilitates the specific parametrisation of each dataset allowing both narrow or broad peak calling, as well as comparisons between the conditions using multiple statistical settings. Finally, complete reports are produced at the end of the bioinformatic and the statistical part of the analysis, which facilitate the data quality control and the interpretation of the results.

We explored the discriminative power of the statistical settings for the differential analysis, using a published dataset of three histone marks (H3K4me3, H3K27ac and H3K4me1) and two transcription factors (Oct4 and Klf4) profiled with ChIP-seq in two biological conditions (shControl and shUbc9). We show that distinct results are obtained depending on the sources of ChIP-seq variability and the dynamics of the chromatin factor under study. We propose that ChIPflow can be used to measure the richness of the epigenomic landscape underlying a biological process by identifying diverse regulatory regimes and the associated genes sets.

## Introduction

Best practices for ChIP-seq analysis have been defined and validated for a long time by the ENCODE consortium [2, 15]. Nevertheless, among the many available tools for ChIP-seq analysis most do not include key data processing steps and statistical controls necessary for the robust and reproducible description of the local epigenomic landscape. Such is the case of the cross-correlation calculation to assess the quality of the immunoprecipitation and the Irreproducible Discovery Rate (IDR) [15, 18] for the estimation of replicate reproducibility, whose automation has proven to be difficult. In contrast, standards for the differential analysis of RNA-seq datasets have been widely studied and implemented, so that users can do an informed choice of the method that will optimise sensitivity and sensibility [8, 17]. This is not the case for ChIP-seq, where most researchers are not aware of the effect that a statistical setting can have on their comparisons, depending on the dynamics of the chromatin factor under study.

We present ChIPflow, a pipeline that streamlines the complete process from raw data to peak calling with the identification of reproducible peaks if replicates are available. ChIPflow has multiple running modes: with/without replicates, one or multiple modification/transcription factors (TFs). It is designed to facilitate the analysis when the parameters are known. However, it also permits to efficiently estimate the optimal parameter values by iterating the analysis without repeating time consuming steps such as mapping and deduplication.

ChIPflow can also take care of the down stream analysis to evaluate the differential marking or binding between multiple biological conditions. This implies an additional level of complexity that is unknown for RNA-seq experiments. ChIP-seq experiments are performed for histone modifications with different nucleation, spreading and maintenance kinetics, as well as proteins with variable affinity to tightly or loosely compacted chromatin. As a consequence, the proper quantification of the marking/binding dynamics needs to consider these differences. ChIPflow provides a flexible set of statistical settings with multiple options to adjust the quantification and subsequent comparison. These include: linear, nonlinear and spike-in methods for the library size normalisation as well as two different variance estimation approaches, limma [27] and DESeq2 [22].

As a proof of concept, we apply the various statistical settings proposed by ChIPflow on a published dataset [9] with three histone modifications (H3K4me3, H3K27ac and H3K4me1) and two TFs (Oct4 and Klf4) profiled with ChIP-seq and we identified differentially marked/bound peaks between two biological conditions (shControl and shUbc9). We show that the choice of normalisation method and variance estimation model influences the differential analysis in a modification/TF dependent manner. Furthermore we propose that such dependency can be described in terms of the two sources of ChIP-seq variability: one related to the enrichment or read coverage and the other connected to the position of maximum enrichment or summit. For a stable modification like H3K4me3 with low enrichment and position variability, distinct sets of differentially marked peaks are identified depending on the statistical setting. Instead, for a TF characterised by high position variability like Klf4, the majority of differentially bound peaks (∼75% of the total differential analysis results) are found using only limma and a nonlinear normalisation. Most importantly, genes neighbouring those differentially marked/bound peaks present different mean expression changes, suggesting variable transcriptomic regimes and thus potentially diverse regulatory mechanisms.

## Results

### The workflow

ChIPflow is a flexible pipeline designed to deal with various chromatin factors (e.g. histone modifications, TFs) profiled in multiple biological conditions with/without replicates. It is built in five modules that execute specific and interdependent tasks (Figure 1A).

**Figure 1.**
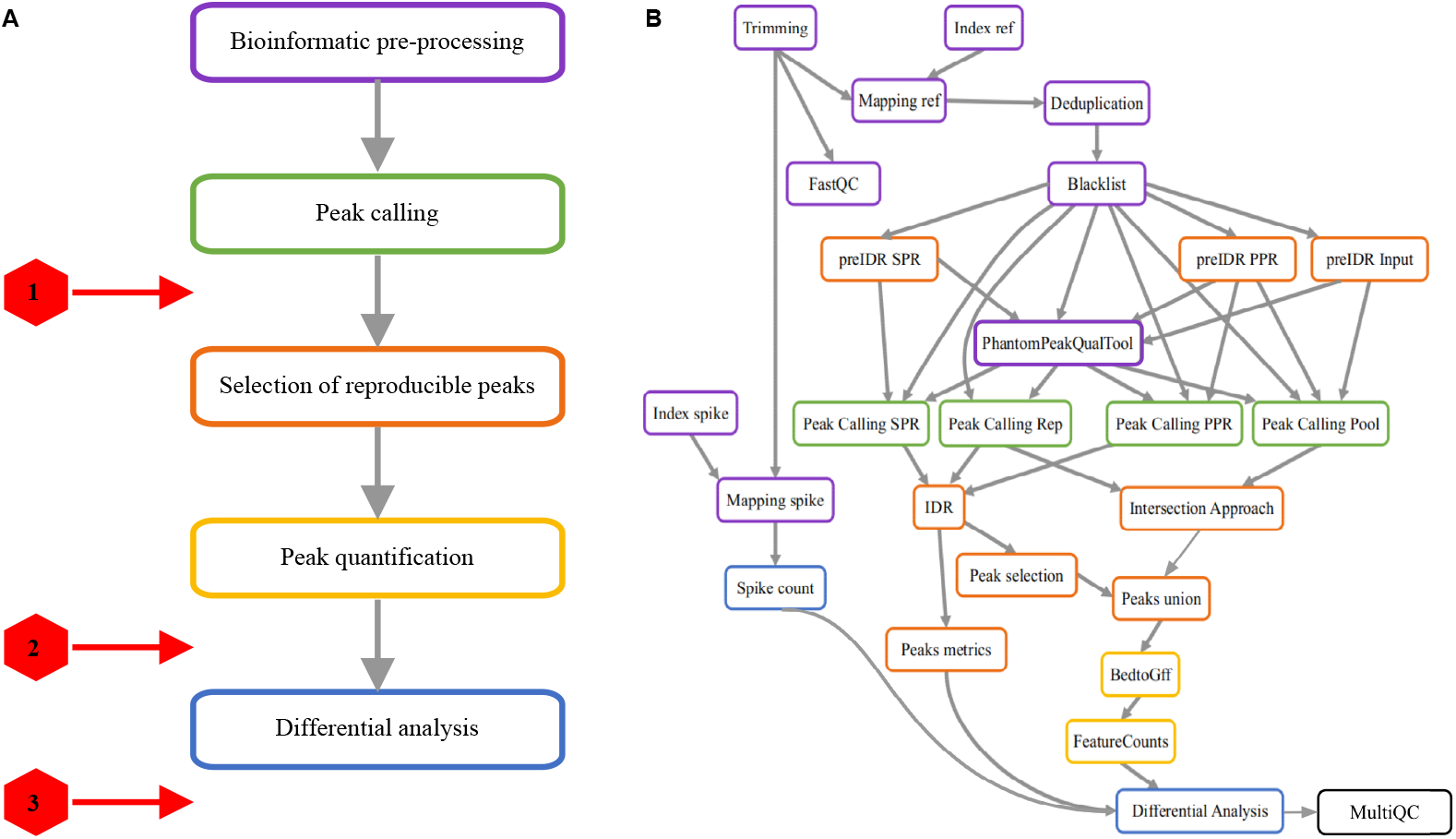
The ChIPflow workflow. **A**. Five ChIPflow modules executing specific and interdependent tasks. Stop signs indicate where the analysis ends, depending on the data provided by the user. Stop 1 if no replicates are available. Stop 2 for datasets with replicates for only one condition. Stop 3 when replicates are available for two or more conditions. **B**. ChIPflow Snakemake rule graph illustrating the input/output dependencies between steps. Border colour indicates the module membership of each rule.

The *bioinformatic pre-processing* module aims at obtaining the deduplicated aligned reads for the peak calling step. It includes: filtering, trimming and quality assessment of the raw data; mapping to the reference genome, correction of PCR amplification biases and removal of blacklisted regions that are the result of artefactual high coverage. For experiments were global changes of histone modifications take place and a spike-in control is added, this module executes the same steps for the exogenous chromatin on the corresponding genome.

Next, the *selection of reproducible peaks* and *peak calling* modules are two intermingled routines that provide the list of top ranking peaks. The peaks that are reproducible between replicates are identified using the Irreproducible Discovery Rate (IDR) or the intersection approach for narrow and broad peaks respectively.

The *peak quantification* module produces the input for the differential analysis, namely the count matrix containing the enrichment quantification (i.e. the number of reads) of each sample over every peak. Rows of this count matrix correspond to the non-redundant set of reproducible peaks merged among all conditions. This guarantees that the quantitative comparisons are performed between genomic regions of equal length.

Finally, the *differential analysis* module is designed to estimate the significance of the observed changes in marking/binding between the user-defined conditions for each peak in the non-redundant set of reproducible peaks. This module is intended to provide a wide range of statistical settings to obtain a comprehensive quantitative description of the epigenomic-mediated process. Given a proper experimental design, without confounding factors, this module permits to correct unwanted technical variability (i.e. batch effects) and to account for systematic technical variability such as different library sizes. It includes linear and nonlinear methods for normalisation and different variance estimation models as implemented in the DESeq2 and limma packages. For experiments with spike-in control this module performs a linear normalisation using the scaling factor calculated from the mapping coverage of the exogenous chromatin (Figure 6).

### ChIPflow different running modes

ChIPflow proposes two main running modes, depending on the specific needs of the user: (i) a production mode for users who want to analyse their datasets using standard, predefined parameters for the peak calling (narrow vs broad peaks) and the differential analysis (library size normalisation method and variance estimation approach); and (ii) an exploratory mode for users dealing with epigenomic projects where the behaviour of the chromatin factor under study is uncertain.

Depending on the specific question or the available data, ChIPflow allows for an analysis at different levels, as illustrated in Figure 1A. A basic analysis stops at the peak calling step when replicates could not be obtained. The outcome in this case will be a list of peaks that can be ranked according to the adjusted p-value or the log fold change of the observed enrichment. Else, if biological replicates are available, the pipeline assess the concordance of peak calls between replicates using the IDR procedure and output a list of reproducible peaks under a given IDR threshold. Finally, for complete experimental designs with multiple replicated biological conditions, ChIPflow performs the differential analysis as defined by the user.

The exploratory mode of ChIPflow allows to test multiple combinations of peak calling and differential analysis parameters avoiding the re-calculation of intermediate steps. The user can therefore gain a complete understanding of the system while minimising the computing time. Some practical examples of the exploratory mode are: calling narrow and broad peaks on the same pre-processed mapping files or comparing normalisation methods for the differential analysis on the same count matrix.

### The importance of providing multiple differential analysis settings

Unlike RNA-seq reads that measure the transcriptional output of a given locus, ChIP-seq reads represent both the probability of marking/binding and its spatial distribution at a given genomic region, i.e. the peak. It follows that ChIP-seq samples have two sources of variability: one related to the enrichment or read coverage of the peak, and the other to the position of the maximum enrichment or summit. In spite of this, the differential analysis of marking/binding is traditionally performed using the methods developed for RNA-seq like DESeq2 and limma.

We analysed a published dataset of three histone modifications (H3K4me3, H3K27ac and H3K4me1) and two TFs (Oct4 and Klf4) profiled using ChIP-seq in two biological conditions (shControl and shUbc9-treated reprogrammable mouse embryonic fibroblasts) [9]. Using ChIPflow, we explored the discriminative power of various statistical settings for the differential analysis and their correlation to the above mentioned sources of ChIP-seq variability.

For each reproducible peak obtained with the IDR procedure, we determined the enrichment per replicate by counting the number of reads overlapping the peak coordinates. We also measured the summit instability of each reproducible peak per replicate. It was defined as the distance between the summit position estimated by pooling IP replicates and the summit of the overlapping peak called by using each IP replicate (Figure 2).

**Figure 2.**
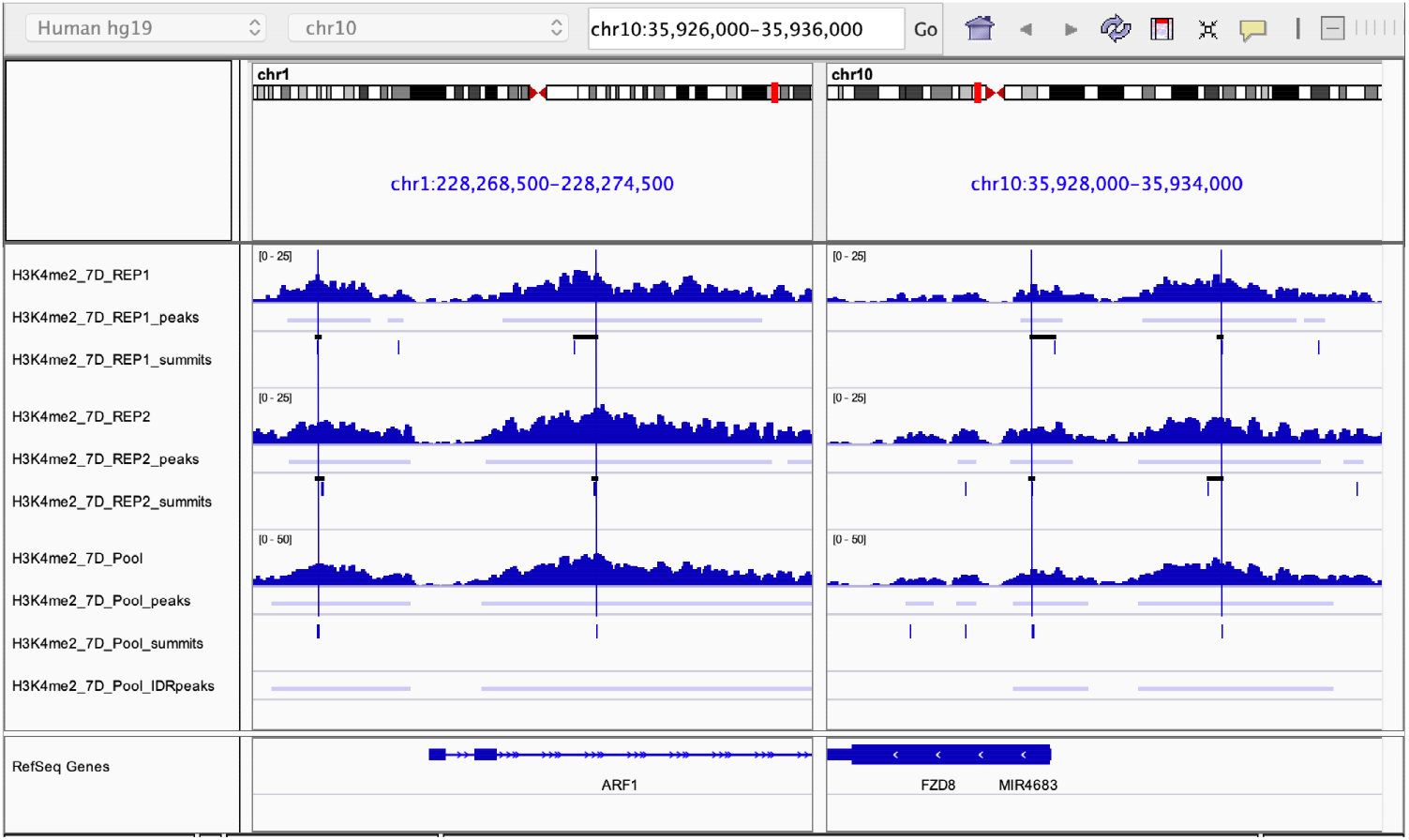
ChIP-seq position variability. Summit instability is defined as the distance between the summit of the peak called in each replicate and the corresponding reproducible peak. Tracks show the IP coverage, peak and summit position for the two replicates separately (top) and pooled (bottom) in two genomic different regions. Vertical blue lines indicate the position of the reproducible peak summit and horizontal black lines the distance between to the corresponding summit in each replicate.

The distribution of read coverage and summit instability for all reproducible peaks is plotted in Figure 3A. Each graph summarises the enrichment and position variability between replicates and biological conditions for a given modification/TF.

**Figure 3.**
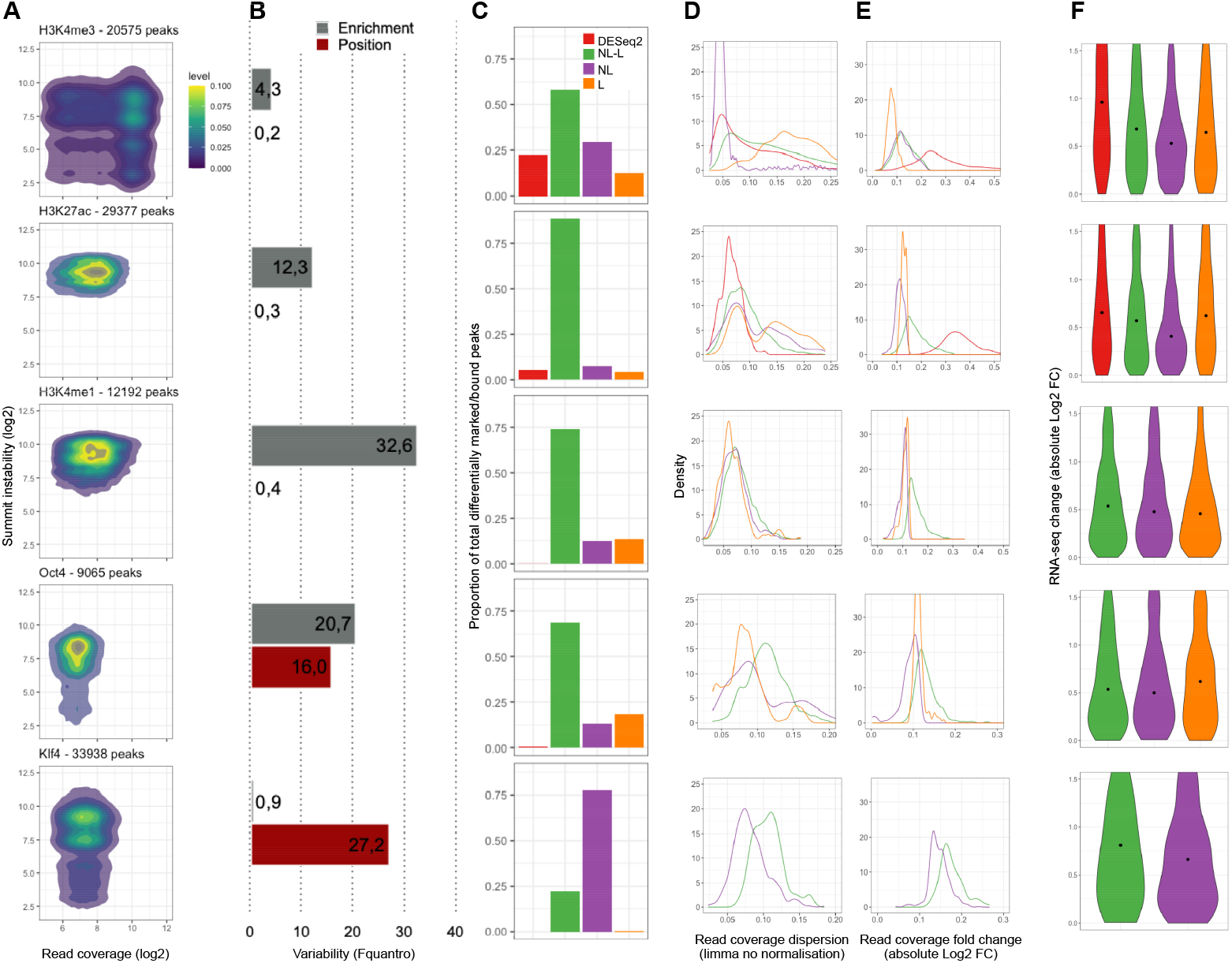
Comparison of the statistical settings for differential analysis. **A**. Kernel density estimate of read coverage and summit instability for all reproducible peaks across replicates of the two biological conditions under study. **B**. Enrichment and position variability estimation using the Fquantro statistic [12] for read coverage and summit instability. **C**. Proportion of total differentially marked/bound peaks obtained using each statistical setting. DESeq2 = DESeq2 with geometric mean normalisation; NL-L = limma with nonlinear and with linear normalisation; NL = limma with nonlinear normalisation only; L = limma with linear normalisation only. **D. E**. Quantitative characterisation of differentially marked/bound peak populations obtained using each statistical setting. Distribution of ChIP-seq read counts dispersion as estimated by limma (D). Distribution of ChIP-seq read counts absolute changes between shUbc9 and shControl (E). Colours correspond to panel C. **F**. Expression dynamics of genes neighbouring differentially marked/bound peak populations. Distribution of RNA-seq read counts absolute changes between shUbc9 and shControl. Colours correspond to panel C.

To quantify the enrichment and position variability of each modification/TF, we calculated the Fquantro statistic for read coverage and summit instability (Figure 3B). This data-driven statistic is a ratio of the mean squared error between-conditions and the mean squared error within-conditions [12].

We performed the differential analysis between shControl and shUbc9 for each modification/TF. We determined the proportion of differentially marked/bound peaks identified using the statistical settings available in ChIPflow: DESeq2 with linear normalisation (geometric mean); limma with linear (scalar) or nonlinear normalisation (quantile or cyclic loess) (Figure 3C).

Depending on the modification/TF, DESeq2 permits the identification of 3 to 25% of the total differentially marked/bound peaks, which are also found by limma independently from the normalisation method. The size of this set of differentially marked/bound peaks diminishes as the enrichment and position variability between conditions increases for the different modifications/TFs (Figure 3B and 3C). This result is compatible with the fact that those peaks have the lowest enrichment dispersion (i.e. variability between replicates) as estimated by limma (Figure 3D). Additionally this is also in agreement with the observation that DESeq2 is a more conservative approach than limma when applied to RNA-seq differential analysis [21].

For the results obtained using limma, three main classes of modifications/TFs can be described (Figure 3B and 3C):

- Stable modification (H3K4me3) with low enrichment and position variability between conditions. Significant marking or binding differences depend on the statistical setting, in particular on whether a linear or nonlinear normalisation method is used.
- Modifications (H3K27ac and H3K4me1) with increasing enrichment variability between conditions and TF (Oct4) with medium variability for both enrichment and position. Differential analysis results are mostly independent from the statistical setting but an additional set of differentially marked/bound peaks (12 to 35% of the total) can be obtained by using specifically a linear or nonlinear normalisation method.
- TF (Klf4) with high position variability and no enrichment variability. More than 75% of the significant differences are detected with a single statistical setting, in this case limma with nonlinear normalisation.

Not surprisingly, there is not an univocal relationship between the number of differentially marked/bound peaks and their quantitative characteristics like the read coverage dispersion or fold change between conditions (Figure 3D and 3E). The largest peak population does not always show the lowest dispersion. Moreover, depending on the modification/TF, a given statistical setting like limma with linear normalisation can identify peak sets with low (H3K4me1 or Oct4) or high (H3K4me3) dispersion (3D). The sole exception is the peak population identified by the DESeq2 approach that, as mentioned above, always includes peaks with low enrichment dispersion (3D).

Interestingly, these peak populations with different quantitative characteristics correspond to distinct expression changes of the neighbouring genes (Figure 3F). This results underlies the biological interest of considering the whole range of differential results when studying epigenomic dynamics.

Our observations illustrate how the use of multiple statistical settings for the differential analysis of marking/binding provides a quantitative description of the various changes that an chromatin factor can undergo and ultimately inform about its regulation mode.

## Discussion

### ChIPflow in comparison to available tools

Since the first publication of the ChIP-seq protocol [3,13] a plethora of methods have been developed and implemented in order to: (i) identify genomic regions with a significant enrichment for the profiled chromatin factor, i.e. peak calling, and (ii) compare the enrichment between biological conditions, i.e. differential analysis.

There is one broad category of methods dedicated to the differential analysis step: DiffBind [32], ChIPQC [33], ChIPComp [7], DBChip [19], MMDiff [28], MAnorm [29]. They mostly differ on the normalisation strategy used to account for systematic technical variability and the statistical test chosen to evaluate the significance of the difference in enrichment per region. Another category of methods perform peak calling and differential analysis: SICER2 [35], MACS2 [37], HOMER [11], RSEG [31], but only diffReps [30], MultiGPS [23] and PePr [36] can take into account replicates for the differential analysis (Figure 4).

**Figure 4.**
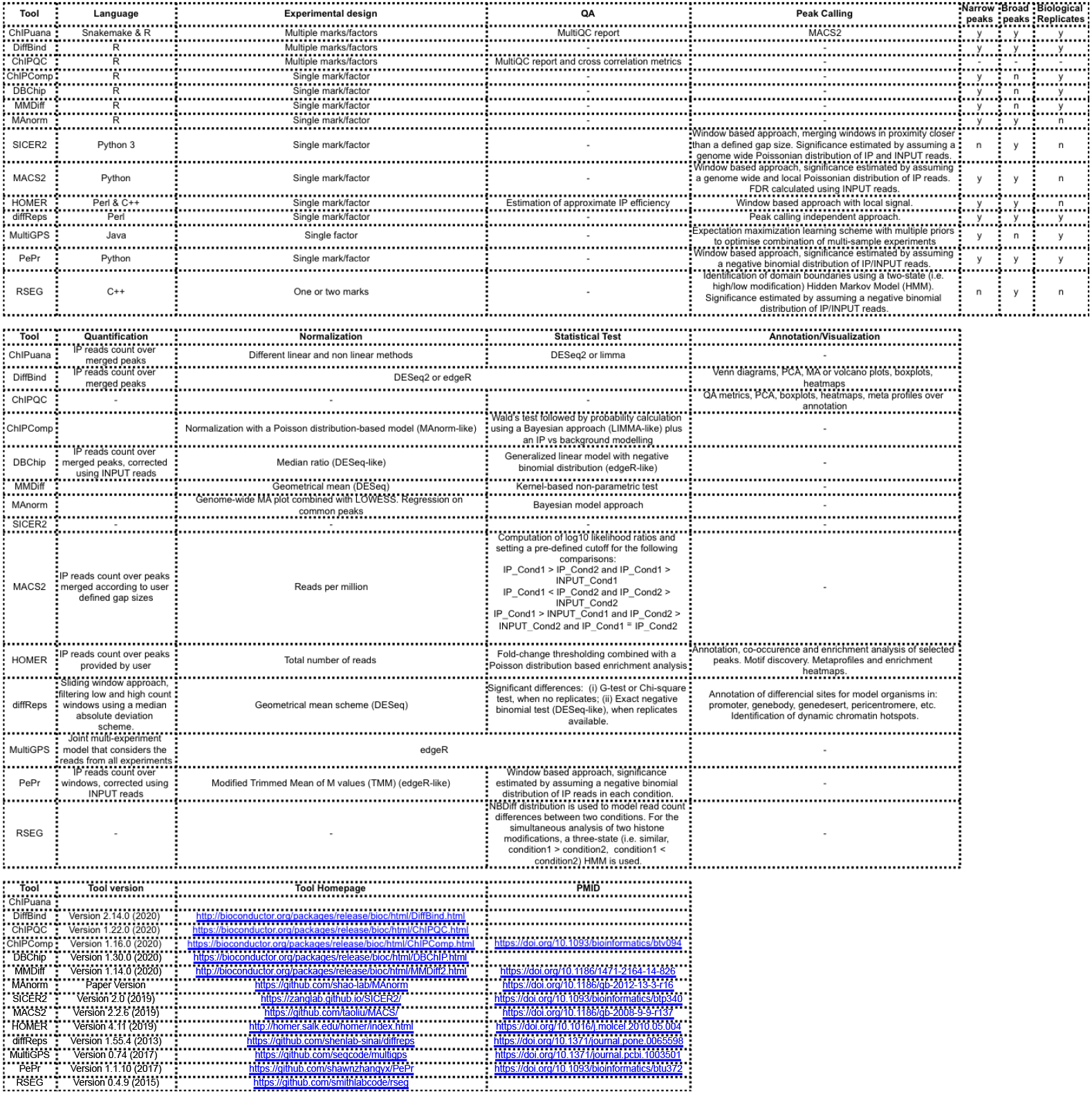
Comparison of available tools for ChIP-seq analysis.

More recently, ChIP-seq pipelines that take care of the intermediate steps from the raw data to the peak calling have been implemented. The GenePipes pipeline [5] includes the calculation of quality control metrics like cross-correlation and sequence bias as well as the functional characterisation of peaks according to their genomic localisation or enriched sequence motifs. However it does not cover the differential analysis. The snakePipes ChIP-seq pipeline [4] does not perform any quality control step but does the differential analysis with the R package CSAW. The nextflow pipeline [25] comprises both quality control metrics calculation and differential analysis using the R package DESeq2 and can be applied on narrow and broad peaks.

To our knowledge, there is no current pipeline that, like ChIPflow, can perform all the following tasks: (i) peak calling starting from the raw sequencing data for both narrow and broad peaks (ii) identification of reproducible peaks between replicates (iii) differential analysis using multiple statistical settings. Of particular interest is the automating of the Irreproducible Discovery Rate (IDR) procedure for the selection of reproducible narrow peaks, which is not implemented by any of the above mentioned methods/pipelines.

The pipeline can process multiple modifications/TFs and conditions simultaneously, with or without replicates. Since every modification/TF is treated separately and according to its experimental design, each one can have a different number of replicates and can be compared between different biological conditions. Last but not least, ChIPflow produces (i) a customised MultiQC that includes quality reports of all the pre-processing steps for all datasets and (ii) a complete report with the results of the differential analysis.

### Beyond a ChIP-seq analysis pipeline

On top of being a complete ChIP-seq analysis pipeline, ChIPflow is a flexible tool to systematically explore the epigenomic landscape underlying a dynamic biological system. It permits the exhaustive exploration of the marking/binding changes that follow or drive from/a perturbation (e.g. mutation, treatment, pathogen infection) of a steady or control state.

Indeed, the various statistical settings of the differential analysis module enable a quantitative estimation of the heterogeneity of the regulatory response. The distinct populations of differentially marked/bound peaks can be interpreted as groups of genomic regions with differing regulatory roles, e.g. related to leaky or not synchronised transcriptional events, or to changes occurring only in a cell sub-population.

Our results on the statistical settings used for the differential analysis of three histone modifications and two TF exemplify the complex relation between the sources of ChIP-seq variation and the ability to identify significant marking/binding changes. Assuming that unwanted technical variation is corrected or accounted for, the ChIP-seq profiling is measuring a dynamic behaviour resulting from the interplay between the intrinsic regulatory characteristics of a chromatin factor and the genomic/nuclear/cellular environment shaped by the biological process under study. These intrinsic factors may include, among others, the writing/reading/erasing kinetics of the histone modifications or the binding affinity of the TFs including their ability to target compacted chromatin. While the sole epigenomic profiling is not enough to deconvolute all those intrinsic and environmental aspects, it can give a sense of the variety of regulatory possibilities. Considering all the above, ChIPflow can be used to measure the richness of the epigenomic landscape and identify potentially diverse regulatory regimes as well as the associated gene sets.

## Methods

ChIPflow is developed in Snakemake [14] and follows the good practices of reproducibility described by the Snakemake’s authors. The workflow is stored in a dedicated git repository https://gitlab.pasteur.fr/hub/chipflow. The main pipeline is in the Snakefile, and all the rules are stored in the workflow/ directory. One rule corresponds to one software or one step and we can split the pipeline into five modules (Figure 1A):

- Bioinformatic pre-processing: all samples are treated independently according to the experimental design.
- Peak calling: IP and INPUT are matched for each modification/TF and condition.
- Selection of reproducible peaks: IPs are evaluated by replicates, self pseudo-replicates and pooled pseudo-replicates for each modification/TF and condition.
- Peak quantification: reproducible peaks of all conditions are merged for each modification/TF and read counts are computed per IP sample.
- Differential analysis: performed per modification/TF.

The pipeline includes a combination of python and Snakemake code. It starts by reading the configuration and design files. Then, it verifies the agreement between the information in the configuration and design files and the raw data file names: in case of discordant information or if any file is missing the pipeline stops immediately. Finally, the design file is analysed to determine which module(s) can be run and where the pipeline will end. From the design file a list of wildcards is built and will be used to automatically allocate each input file to its corresponding Snakemake rule.

For example, if the user chooses to perform the differential analysis but replicates are not provided, the pipeline will automatically stop after the peak calling step. Seemingly, a minimum of two replicates by IP and one INPUT are required to obtain reproducible peaks; at least 2 conditions per modification/TF and reproducible peaks are required to perform the differential analysis (Figure 1A). Since ChIPflow is processing each modification/TF independently, it can run until the reproducible peaks selection for one dataset, and still run the differential analysis module for another one.

### Bioinformatic pre-processing module

The bioinformatic pre-processing module includes quality control of reads, adapters’ trimming, mapping, read deduplication, removal of blacklisted regions, if possible, and calculation of cross-correlation quality metrics.

The tools used within this module are (Figure 1B):

- FastQC for quality control of samples [1].
- Cutadapt for filtering reads and trimming adapters [24].
- bowtie2 for read mapping [16]. Mapping is performed against the IP reference genome, and is also done against the control reference genome for spike-in samples. For user-provided genome sequences, a dedicated rule handles the genome indexing.
- MarkDuplicates from the Picard tools suite for removal of duplicated reads [6].
- intersectBed from the bedtools suite [26] for removal of blacklisted regions. This rule is optional, because blacklisted regions have not been identified for many non-model organisms.
- PhantomPeakQualTools for calculation of cross-correlation metrics [15]. The main script is written in R and the version used in ChIPflow is stored in the workflow/scripts/ directory.

For samples without replicates, the pipeline stops at this point (Figure 1A).

### Peak calling module

After the pre-processing module, enriched regions are identified with the MACS2 peak caller [37]. MACS2 can be run using either narrow or broad mode depending on the expected size of the binding site or the histone marking pattern.

Two inputs are required for the *macs2* rule: a BAM file corresponding to the IP and one corresponding to the INPUT. IP and INPUT BAM files are paired for each modification/TF and condition, according to the design file. Peaks can only be called on an IP with a matched INPUT. If several INPUT BAM files are provided corresponding to different replicates, the first one is used for all IP replicates. This approach assumes that there is no significant variability between INPUT replicates. Optionally, the shift file from PhantomPeakQualTools can be additionally provided if the user prefers to keep the fragment estimation obtained from the cross-correlation calculation. This is particularly useful when MACS2 fails to calculate the fragment size using the default model approach.

Output files are stored in directory with explicit names specifying the peak calling mode (narrow or broad) and the fragment size estimation method (model or no model). This allows the user to run this module several times with different parameters while avoiding to over-write the results.

### Selection of reproducible peaks

The next step is to select the reproducible peaks between biological replicates. The protocol for this depends on the peak calling mode. As a default, the Irreproducible Discovery Rate (IDR) procedure is used for narrow peaks and the intersection approach for broad peaks.

The automated IDR procedure of ChIPflow corresponds to the one described by [15]. It starts with a pre-computing step (Figure 1B) and provides a set of IDR metrics (Figure 5) that are used for the assessment of the replicates’ reproducibility and the final selection of reproducible peaks.

**Figure 5.**
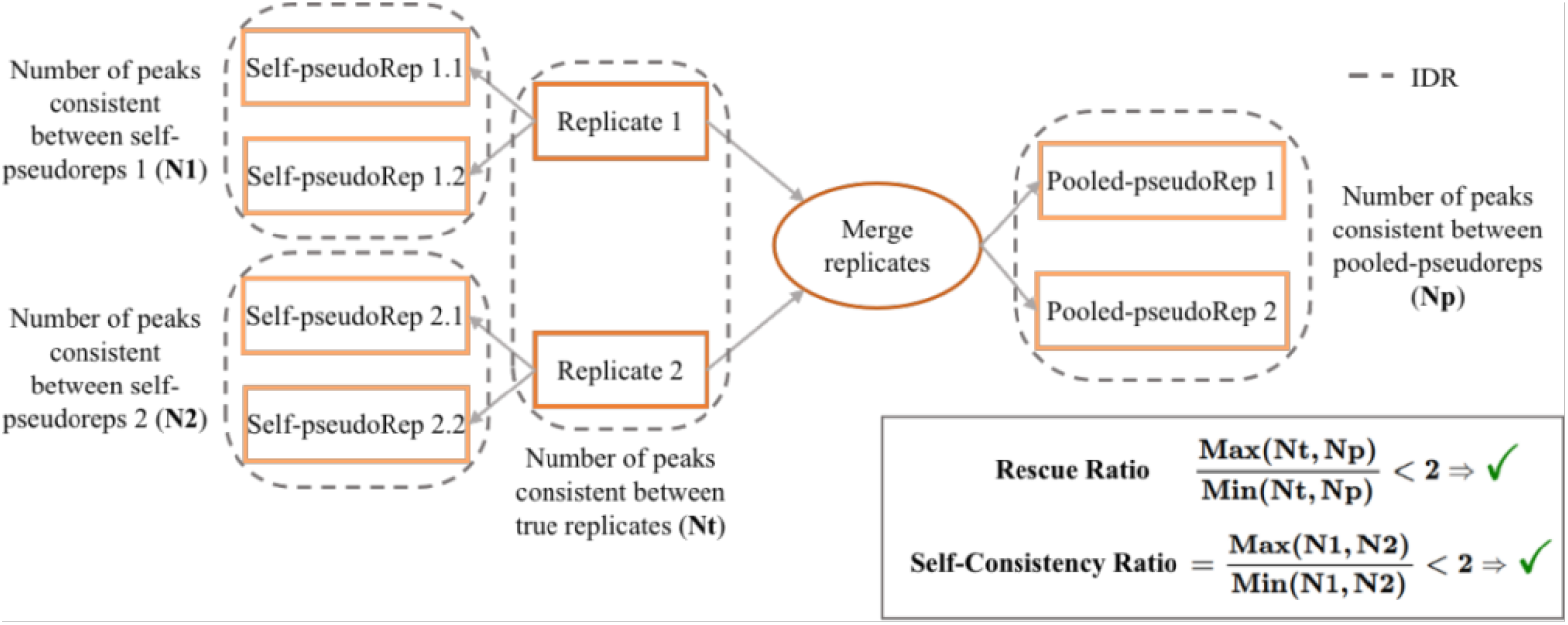
Irreproducible Discovery Rate (IDR) procedure as described in [15]. Steps of the IDR are represented inside dashed line boxes. IDR metrics used for the assessment of the replicates’ reproducibility are shown inside the continuous line box.

To compute the IDR per peak multiple files have to be pre-computed. Those files are produced by the pre-IDR submodule as defined below:

- *preIDR_SPR* rule generates two IP self pseudo-replicates for each IP replicate by randomly splitting the mapped reads into two BAM files.
- *preIDR_PPR* rule generates two IP pooled pseudo-replicates by merging all mapped IP reads in a single BAM and then randomly splitting it into two BAM files.
- *preIDR_Pool* rule generates, if INPUT replicates are provided, a pooled INPUT BAM file.

This step takes place before the peak calling. Peaks are called on those seven BAM files, with the INPUT as control except for the pooled IP for which the pooled INPUT is used as control if available.

Then the IDR score is estimated for each peak of the IP replicates, self pseudo-replicates and pooled pseudo-replicates. The rescue ratio and self-consistency ratio are calculated as described by [15]. Finally, the reproducible peaks are selected among the peaks called on the pooled IPs and having the lowest IDR score.

The IDR procedure is optimised for narrow peaks. For this reason we included the intersection approach to choose reproducible broad peaks. First, peaks are filtered by selecting those with a q-value lower than 0.01 and a log fold change higher than 3. Second, the proportion of overlap per peak between replicates is determined using intersectBed (from BEDtools suite). Only peaks with an overlap of at least 80% of the longest peak length are kept. The user can force the use of the intersection approach for narrow peaks.

### Differential analysis

Differential analysis is performed for modifications/TFs matching the following requirements:

- A minimum of two biological conditions.
- Two replicates for each condition from which reproducible peaks have been selected.

A union peak file is obtained by merging the reproducible peaks coordinates for all the conditions of one modification using mergeBed from BEDtools with default parameters. Merged peak summit is calculated as the mean of all merged peak summit positions.

The count matrix is then generated for all IP BAM files using featureCounts [20] and the peak coordinates in the union peak file.

The differential analysis is performed to detect differentially marked/bound peaks between the biological conditions. For this, we adapted the SARTools R package [34] and created chIPflowR which is freely available at https://gitlab.pasteur.fr/hub/chipflowr. Two approaches to model variability (DESeq2 and limma) as well as linear and nonlinear normalisation methods (scalar, quantile, cyclic loess, spike-in) are available to perform the differential analysis (Figure 6). Using the count matrix generated by featureCounts, chIPflowR carries out the quality assessment of the data using dedicated plots, runs the differential analysis and exports (i) a Rmarkdown HTML report describing the analysis process and (ii) the tables containing the differentially marked/bound peaks coordinates and corresponding statistics.

**Figure 6.**
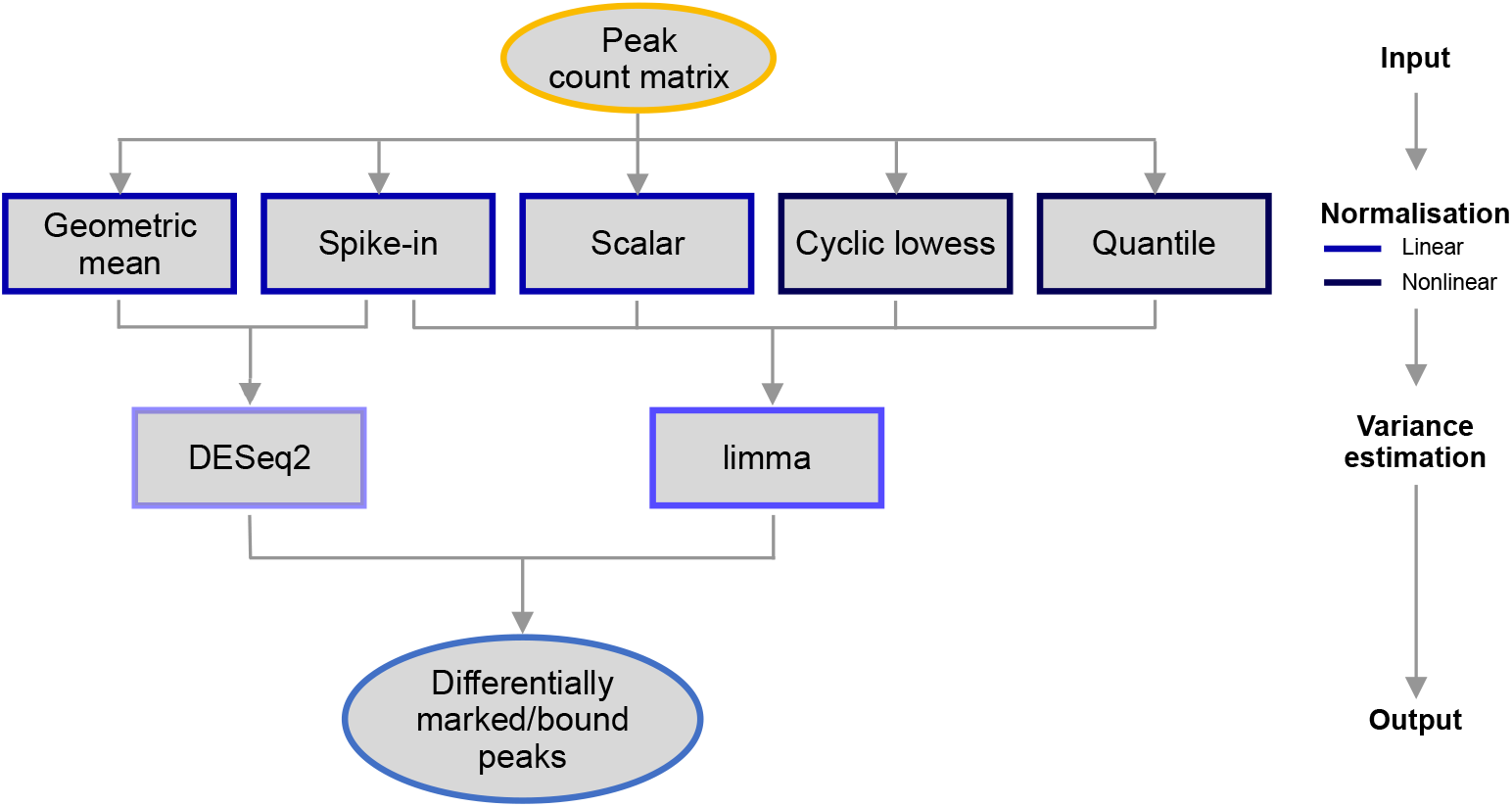
ChIPflow differential analysis module with the statistical settings implemented in chIPflowR. Possible combinations of normalisation method and variance estimation approach are shown.

### Analysis report

Finally, a MultiQC [10] report is compiled, which concatenates all relevant information and metrics from the log files produced during the pipeline execution. It provides a global view of the results (Figure 7).

**Figure 7.**
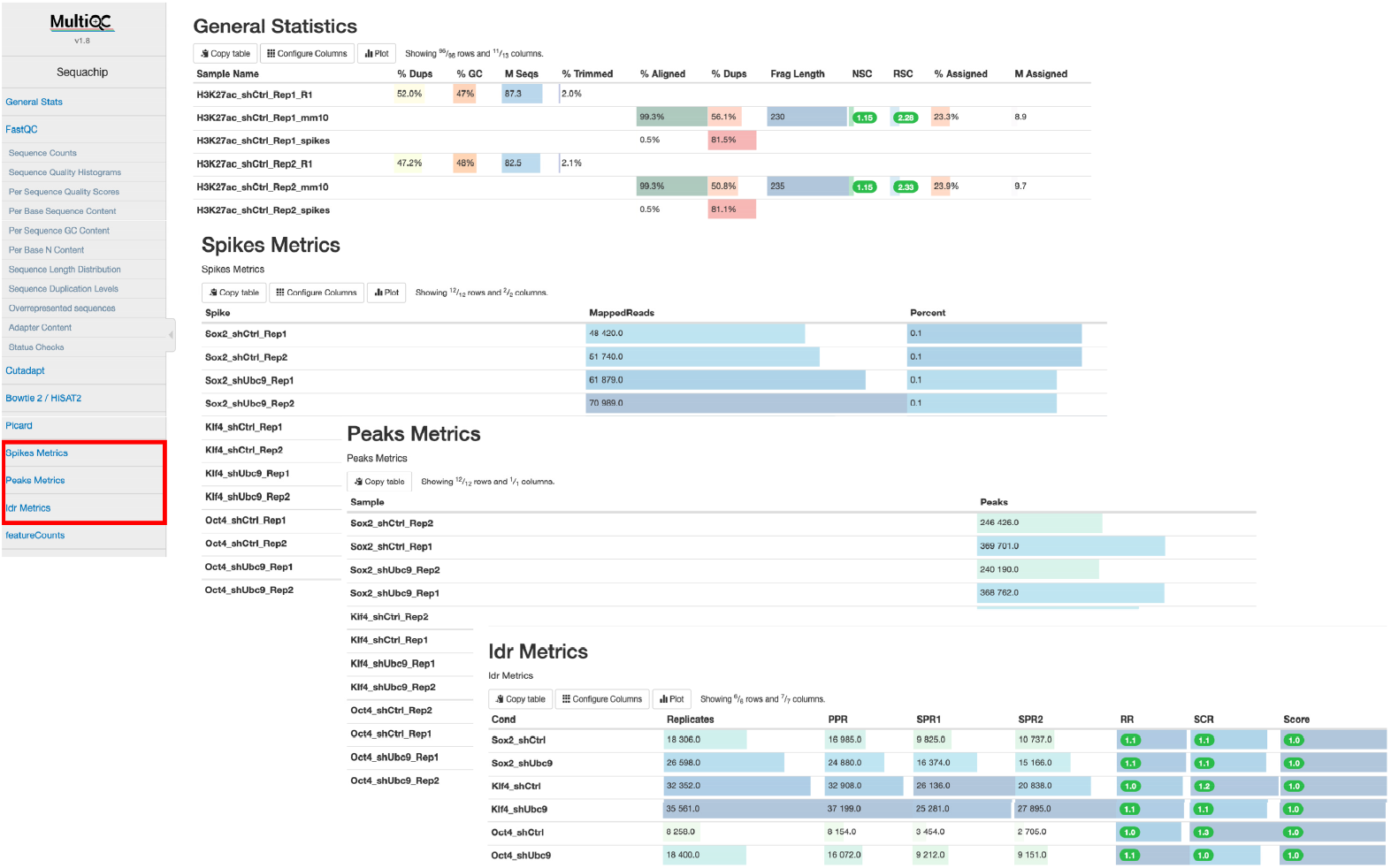
Customised MultiQC report. All classical MultiQC modules are included to provide an interactive report of every step in the analysis. Modules that were added to adapt the report for ChIP-seq datasets are highlighted in red and presented as partial screenshots.

A particular effort was put in the customisation of the MultiQC config file aiming at offering a complete and interactive final report. In addition to the classic MultiQC modules such as bowtie2 or picard-tools, we added homemade chunks: a table summarising the number of peaks called per IP dataset; for spike-in datasets, a table containing the mapping statistics on the exogenous genome. Moreover, a whole section is dedicated to the IDR results. Rescue Ratio, Self-Consistency Ratio and IDR score are tabulated for replicates, self pseudo-replicates and pooled pseudo-replicates, facilitating the identification of reproducible replicates. To further simplify the quality assessment, samples that failed or succeeded according to the PhantomPeakQual tools and the IDR metrics are highlighted using a MultiQC formatting rule.

### Distribution

The pipeline is freely available at https://gitlab.pasteur.fr/hub/chipflow/.

## Author Contributions

MD and CC performed the dataset analysis. RL and MD conceived and wrote the Snakemake pipeline. MD and HV created the chIPflowR R package. HV gave input in the statistical analysis. RL and AP debugged and tested the pipeline. CC conceived and supervised the study. CC, MD, RL wrote the paper with input from all the authors.

## Acknowledgments

Authors would like to thank Thomas Cokelaer for his early input, Thomas Bigot and Blaise Li for helping on the Snakemake development and the singularity packaging, Julien Guglielmini for his advice on Bash and all members of the Hub of Bioinformatics and Biostatistics for useful discussions.

